# Rapid diagnosis of SCA36 in a three-generation family using short-read whole genome sequencing data

**DOI:** 10.1101/851675

**Authors:** Haloom Rafehi, David J. Szmulewicz, Kate Pope, Mathew Wallis, John Christodoulou, Susan M White, Martin B Delatycki, Paul J Lockhart, Melanie Bahlo

## Abstract

**Background:** Spinocerebellar ataxias (SCA) are often caused by expansions of short tandem repeats (STRs). Recent methodological advances have made repeat expansion (RE) detection with whole genome sequencing (WGS) feasible.

**Objectives:** To determine the genetic basis of ataxia in a multigenerational Australian pedigree, with autosomal dominant inheritance.

**Methods and Results:** WGS was performed on three affected relatives. The sequence data was screened for known pathogenic REs using two repeat expansion detection tools: exSTRa and ExpansionHunter. This screen provided a clear and rapid diagnosis (<five days from receiving the sequencing data) of SCA36, a rare form of ataxia caused by an intronic GGCCTG RE in *NOP56*.

**Conclusions:** the that diagnosis of rare ataxias caused by REs is highly feasible and cost effective with WGS. We propose that WGS be implemented as the frontline, cost effective methodology for molecular testing of individuals with a clinical diagnosis of ataxia.

## Introduction

Spinocerebellar ataxias (SCA) comprise a group of rare, progressive neurological disorders that can be caused by deleterious coding-sequence point mutations or pathogenic expansions of short tandem repeats (STRs). Ataxias caused by repeat expansions (REs) of STRs are difficult to diagnose, with low throughput methods such as locus-specific sizing or repeat primed PCR assays being most commonly performed ^1^. Clinical testing is routinely available for SCAs caused by more common REs such as Friedreich’s ataxia, SCA1, 2, 3, 6 and 7, but not rarer forms such as CANVAS, SCA 8, 10, 12, 17, 31, 36 and 37. Hence molecular diagnosis of ataxias due to REs, particularly less common REs such as those underlying SCA36, can be hard to obtain. In addition, current RE tests are expensive and time consuming, sometimes requiring international shipping of biospecimens. This has multiple negative outcomes, including uncertainty and a potential diagnostic odyssey for patients, while clinicians and health providers are unable to offer accurate prognosis, counselling and best clinical care.

Recent methodological advances in bioinformatics have made the simultaneous testing for all known REs through whole genome sequencing (WGS) feasible ^2^. Additionally, multiple novel REs have recently been discovered in neurological disorders using WGS, including in epilepsy ^3-6^ and ataxia ^7-9^. Indeed, there is a growing recognition of the importance of REs and potential contribution to unsolved neurogenetic disease ^10^. This study demonstrated the potential utility of WGS for testing multiple SCA associated loci simultaneously.

In a recent study, we utilised WGS to discover a novel AAGGG RE in the gene replication factor 1 (*RFC1*) that causes cerebellar ataxia with neuropathy and bilateral vestibular areflexia syndrome (CANVAS) ^7^. We provided a genetic CANVAS diagnosis to patients from 18 families in our Australian CANVAS cohort. Additionally, we used the WGS data to re-diagnose patients who tested negative for the *RFC1* RE with other forms of ataxia, including SCA3, also caused by a RE.

Here, we present a case study in which a three generation Australian pedigree with autosomal dominant ataxia was rapidly diagnosed with SCA36 using WGS-based bioinformatic approaches, and subsequently validated by two independent international service providers. SCA36 is a very rare ataxia with slow progression, and is characterized by adult-onset gait and appendicular ataxia, oculomotor abnormalities, cerebellar dysarthria and hyperreflexia. A definite clinical diagnosis of SCA36 is difficult due to overlap with other ataxias, and therefore targeted testing of this locus is not routinely considered. This issue is compounded by the fact that in Australia and many other countries there is no accredited diagnostic testing available for SCA36. A small number of international service providers offer single locus SCA36 testing, but the test is expensive, turnaround times can be long, and cannot provide the exact size of the RE. This study highlights that accredited diagnostic WGS, which is available in many countries, can provide an alternative single test to rapidly diagnose SCA36 and other unsolved ataxias, effectively addressing a previously unmet clinical need.

## Methods

Institutional Ethics Committee approval was provided by the Royal Children’s Hospital (Melbourne, HREC 28907), and written informed consent was obtained from all participants prior to study. Clinical details were collected from clinical assessments and review of medical records. Genomic DNA was isolated from blood using standard techniques and WGS was performed fee for service by the Australian Genome Research Facility. Libraries were prepared using a PCR free kit (TruSeq) and sequenced using a Novaseq (Illumina). Alignment and haplotype calling were performed based on the GATK best practice pipeline. All samples were aligned to the hg19 reference genome using BWA-mem. Duplicate marking, local realignment, and recalibration were performed with GATK. Samples were screened for 12 ataxia STRs: SCA1,2,3,6,7,8,10,12,17,36, Friedreich’s ataxia and CANVAS. RE detection tools exSTRa and ExpansionHunter were used, using default parameters, with additional parameters for ExpansionHunter: inclusion of off-target reads and read-depth of 30 and min-anchor-mapq of 20. Results were compared to 167 control WGS samples from the GTEx project.

## Results

A large Australian family of European ancestry with seventeen individuals affected by ataxia over three generations was recruited to this study (Figure 1). Segregation is consistent with autosomal dominant inheritance, with apparently complete penetrance. At least 18 members of this family in 4 generations present with ataxia. Onset of symptoms is in mid adulthood and is slowly progressive. Lifespan is not significantly reduced with a number of affected individuals living into their 80s. A detailed neurological examination was undertaken on the three subjects and found that all displayed an ataxic gait, characterized by a broadened base of support, lateral veering, reduced walking speed, irregular foot placement, an increased stance phase and increased foot rotation angles ^11,12^. Oculomotor abnormalities were consistent with cerebellar impairment and included saccadic horizontal and vertical visual pursuits ^13^ and vestibulo-ocular reflex suppression ^14^; and dysmetric saccades to target ^15^. Cerebellar dysarthria with consequent reduction in speech intelligibility was also present ^16,17^. Appendicular ataxia was seen in all four limbs and manifested by dysmetria ^18,19^ and intention tremor ^18,20^. Global upper and lower limb hyperreflexia was also present on examination ^21^.

**Figure 1.**
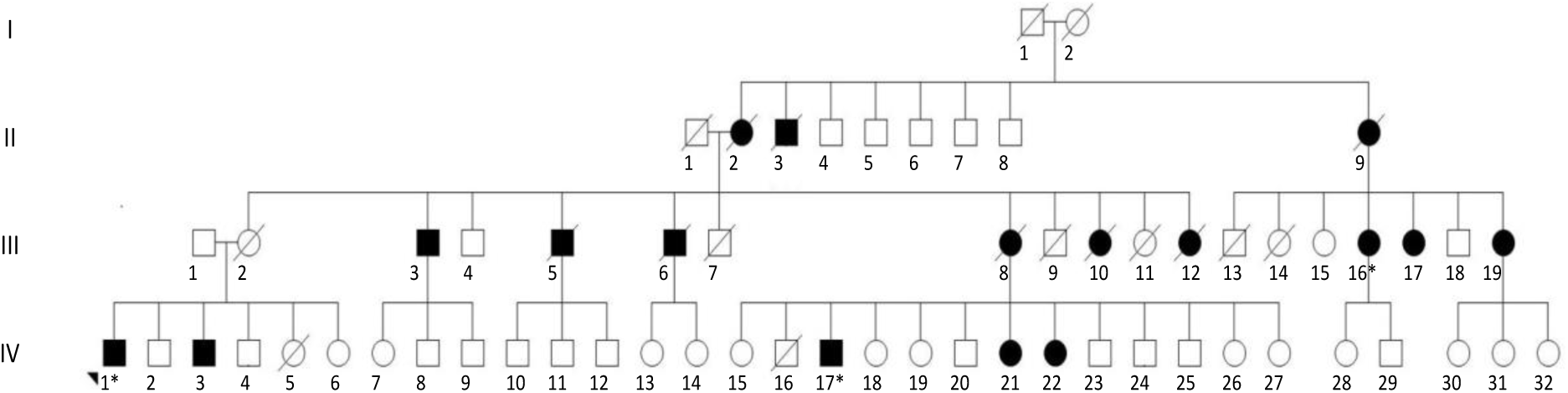
Multigenerational ataxia family pedigree. Individuals with confirmed ataxia have been shaded in black. The individuals with WGS data are indicated by the *.

WGS was performed for three affected individuals from the family (Figure 1) and the data was analysed for a pre-defined list of 12 candidate repeat expansions known to cause ataxia (SCA1,2,3,6,7,8,10,12,17,36, Friedreich’s ataxia and CANVAS) using exSTRa ^22^ and ExpansionHunter ^23^. This analysis identified the SCA36 RE in all three individuals, but none of the 167 controls. Based on evidence from the repeat read distributions plotted by exSTRa the RE was present in a heterozygous state in all affected individuals sequenced (Figure 2A). ExpansionHunter was used to estimate the size of the expanded allele determining a lower bound of ∼800 repeat units (Figure 2B). This is a lower bound due to the limitations of sizing of very large expansions with short-read sequencing, with a known bias towards estimating fewer repeat motifs. This is a significant distinction as the pathogenic threshold for SCA36 has previously been described as greater than 650 repeats ^24^. Individuals with normal repeat size display a motif range of 1-12 repeat units. More precise sizing of SCA36 RE requires a Southern blot, however no such Southern blot is currently available commercially to our knowledge due to the rarity of this RE. Visualisation of the locus in IGV confirms a large RE present in all three individuals (Figure 2C).

**Figure 2.**
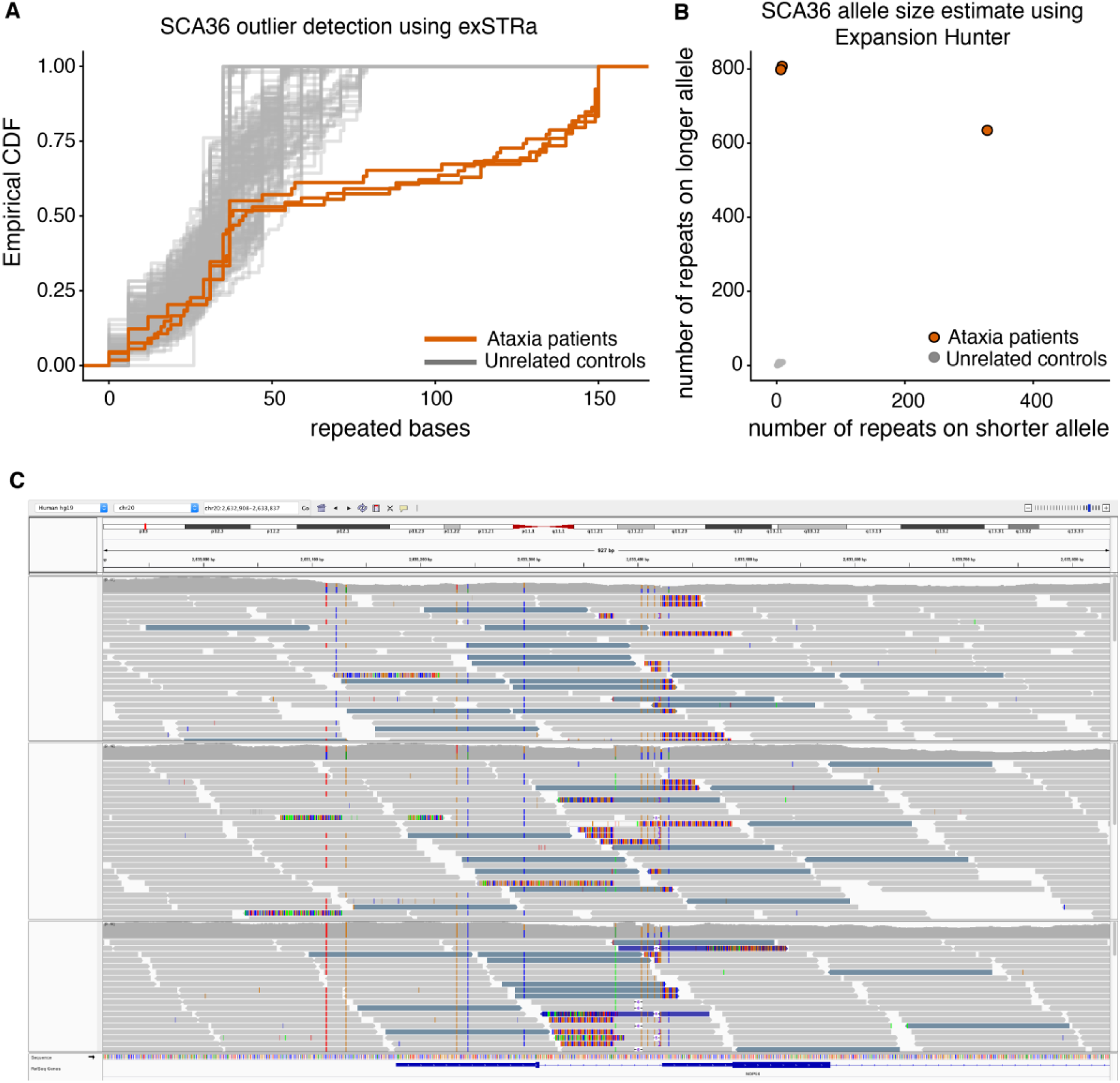
The SCA36 RE was detected using (A) exSTRa and (B) ExpansionHunter. (C) Visualisation in IGV of the GGCCTG RE in the three patients. The blue-grey reads represent reads where the mate has mismapped to a GGCCTG-rich locus (chr5:145,838,637).

Subsequent to identification of SCA36, we performed confirmatory linkage analysis using genotypes extracted from the WGS data with LINKDATAGEN ^25^. While underpowered, this analysis identified 15 regions of identity by descent with LOD scores between 1 and 1.5 (data not shown). One of these regions (chr20:257608-7368304, LOD=1.5024, 20p13) overlapped SCA36, and also encompasses the genes for SCA23 (*PDYN*) and SCA35 (*TGM6*). A second linkage region on chromosome 4 intersects with the gene that causes SCA41. No potentially pathogenic sequence variants or RE were identified in these genes in the three affected individuals.

Clinical examination confirmed features consistent with SCA36. To confirm the diagnosis, molecular diagnostic SCA36 testing was ordered. As diagnostic testing is not currently available in Australia, diagnosis was sought through two independent international service providers (Fulgent and MNG Laboratories). Both returned a positive result, identifying one allele as greater than 70 repeats. However, only alleles >650 are considered full-penetrance and diagnostic of SCA36, while alleles between 15 and 650 repeats are considered variants of uncertain significance. Neither provider offers Southern blot or any other test to accurately size the RE.

## Discussion

Here, we report for the first time that WGS can provide rapid and efficient diagnosis of SCA36. This is of significant clinical importance, given the difficulty and lack of availability of diagnostic testing for SCA36 worldwide, with affected individuals remaining underdiagnosed for this genetic mutation. Importantly, other than Southern blot, which is rarely used, no other test can identify >600 repeats for the SCA36 locus. We demonstrate that WGS can offer a rapid and efficient method for to screen for ataxia-associated REs, such as SCA36.

SCA36 is a rare monogenic form of ataxia that is caused by an intronic GGCCTG hexanucleotide repeat expansion in *NOP56* (OMIM #614153). It was first identified in patients of Japanese ancestry, using linkage mapping in three unrelated families ^24^. Nine unrelated affected individuals were identified in a large Japanese ataxia cohort (n=251). These individuals shared a common haplotype, suggesting a single founder event for SCA36 in the Japanese population. Individuals with SCA36-mediated ataxia have since been identified in China, the US, Italy and Spain, with the latter country demonstrating a relatively high incidence of SCA36, owing to a founder effect ^26^. It appears likely that there are multiple founder events for SCA36 RE as has been observed for other repeat expansions, such as Huntington disease ^27^ and SCA3 ^28^.

SCA36 is exceptionally rare - detected in 3.6% of patients in the initial Japanese study ^24^, and only 0.7% of unsolved ataxia patients in a US study (n=577) ^29^. Therefore, routine screening of ataxia patients for SCA36 using RP-PCR or by Southern blot is neither practical or cost effective. We propose that changing clinical practice to utilise WGS as a stand-alone frontline test is a potentially cost-effective and efficient diagnostic pathway for individuals with a clinical diagnosis of ataxia. This method allows for the screening of all known RE simultaneously while also allowing examination of point mutations in known ataxia genes. The WGS data can subsequently be used for novel gene discovery in unsolved cases.

The rapidity and ease of genetic diagnosis for this family with WGS-based RE detection analysis, after decades of genetic testing, will likely extrapolate to other rare REs as well. In the ataxias more broadly, numerous diagnostic procedures such as MRI, nerve conduction studies and cognitive testing are performed, in part to inform diagnostic genetic testing. However our experience, and globally, is that despite these extensive investigations, a diagnosis is only achieved in ∼30% of cases ^30-32^. The diagnostic journey can be long (up to 20 years) and often unsuccessful. We strongly advocate the implementation of WGS for molecular diagnostic testing of individuals with a clinical diagnosis of ataxia. To facilitate these analyses we maintain up to date lists of known RE for all currently used bioinformatic RE detection tools. These can be obtained from https://github.com/bahlolab/Bio-STR-exSTRa.

## Acknowledgements

We acknowledge contributions from Michael Hildebrand, Natasha Brown and Samuel Berkovic. P.J.L is supported by the Vincent Chiodo Foundation. M.B. was supported by an NHMRC Senior Research Fellowship (GNT1102971). D.J.S is supported by an NHMRC Early Career Fellowship (1129595). Whole genome sequencing costs were funded by Austin Health and the Victorian Medical Research Acceleration Fund through the Austin Health Undiagnosed Diseases Program. This work was supported in part by the Victorian Government’s Operational Infrastructure Support Program and the NHMRC Independent Research Institute Infrastructure Support Scheme (IRIISS).

## Authors’ roles

1. Research project: A. Conception, B. Organization, C. Execution;
2. Statistical Analysis: A. Design, B. Execution, C. Review and Critique;
3. Manuscript: A. Writing of the first draft, B. Review and Critique.

HR: 1A,B,C, 2A,B,C, 3A,B.

DJS: 1B,C, 3B

KP: 1B,C, 3B

MW:1B, 3B

JC: 1B, 3C

SMW: 1B, 3C

MBD: 1A,B,C, 2C, 3B

PJL: 1A,B,C, 2AC, 3B

MB: 1A, 2AC, 3B.

## References

1. de Silva RN, Vallortigara J, Greenfield J, Hunt B, Giunti P, Hadjivassiliou M. Diagnosis and management of progressive ataxia in adults. Pract Neurol 2019;19(3):196–207.

2. Bahlo M, Bennett MF, Degorski P, Tankard RM, Delatycki MB, Lockhart PJ. Recent advances in the detection of repeat expansions with short-read next-generation sequencing. F1000Res 2018;7.

3. Ishiura H, Doi K, Mitsui J, et al. Expansions of intronic TTTCA and TTTTA repeats in benign adult familial myoclonic epilepsy. Nat Genet 2018;50(4):581–590.

4. Corbett MA, Kroes T, Veneziano L, et al. Intronic ATTTC repeat expansions in STARD7 in familial adult myoclonic epilepsy linked to chromosome 2. Nat Commun 2019;10(1):4920.

5. Florian RT, Kraft F, Leitao E, et al. Unstable TTTTA/TTTCA expansions in MARCH6 are associated with Familial Adult Myoclonic Epilepsy type 3. Nat Commun 2019;10(1):4919.

6. Yeetong P, Pongpanich M, Srichomthong C, et al. TTTCA repeat insertions in an intron of YEATS2 in benign adult familial myoclonic epilepsy type 4. Brain 2019;142(11):3360–3366.

7. Rafehi H, Szmulewicz DJ, Bennett MF, et al. Bioinformatics-Based Identification of Expanded Repeats: A Non-reference Intronic Pentamer Expansion in RFC1 Causes CANVAS. Am J Hum Genet 2019;105(1):151–165.

8. Cortese A, Simone R, Sullivan R, et al. Biallelic expansion of an intronic repeat in RFC1 is a common cause of late-onset ataxia. Nat Genet 2019;51(4):649–658.

9. van Kuilenburg ABP, Tarailo-Graovac M, Richmond PA, et al. Glutaminase Deficiency Caused by Short Tandem Repeat Expansion in GLS. N Engl J Med 2019;380(15):1433–1441.

10. Lohmann K, Bruggemann N. Rediscovery of repeat expansions: Solving the unsolved cases. Mov Disord 2019;34(9):1300.

11. Stolze H, Klebe S, Petersen G, et al. Typical features of cerebellar ataxic gait. J Neurol Neurosurg Psychiatry 2002;73(3):310–312.

12. Buckley E, Mazza C, McNeill A. A systematic review of the gait characteristics associated with Cerebellar Ataxia. Gait Posture 2018;60:154–163.

13. Thier P, Ilg UJ. The neural basis of smooth-pursuit eye movements. Curr Opin Neurobiol 2005;15(6):645–652.

14. Chambers BR, Gresty MA. The relationship between disordered pursuit and vestibulo-ocular reflex suppression. J Neurol Neurosurg Psychiatry 1983;46(1):61–66.

15. Sharpe JA, Lo AW, Rabinovitch HE. Control of the saccadic and smooth pursuit systems after cerebral hemidecortication. Brain 1979;102(2):387–403.

16. Ackermann H, Vogel M, Petersen D, Poremba M. Speech deficits in ischaemic cerebellar lesions. J Neurol 1992;239(4):223–227.

17. Lechtenberg R, Gilman S. Speech disorders in cerebellar disease. Ann Neurol 1978;3(4):285–290.

18. Vilis T, Hore J. Central neural mechanisms contributing to cerebellar tremor produced by limb perturbations. J Neurophysiol 1980;43(2):279–291.

19. Manto M. Mechanisms of human cerebellar dysmetria: experimental evidence and current conceptual bases. J Neuroeng Rehabil 2009;6:10.

20. Hore J, Flament D. Evidence that a disordered servo-like mechanism contributes to tremor in movements during cerebellar dysfunction. J Neurophysiol 1986;56(1):123–136.

21. Paulson HL. The spinocerebellar ataxias. J Neuroophthalmol 2009;29(3):227–237.

22. Tankard RM, Bennett MF, Degorski P, Delatycki MB, Lockhart PJ, Bahlo M. Detecting Expansions of Tandem Repeats in Cohorts Sequenced with Short-Read Sequencing Data. Am J Hum Genet 2018;103(6):858–873.

23. Dolzhenko E, van Vugt J, Shaw RJ, et al. Detection of long repeat expansions from PCR-free whole-genome sequence data. Genome Res 2017;27(11):1895–1903.

24. Kobayashi H, Abe K, Matsuura T, et al. Expansion of intronic GGCCTG hexanucleotide repeat in NOP56 causes SCA36, a type of spinocerebellar ataxia accompanied by motor neuron involvement. Am J Hum Genet 2011;89(1):121–130.

25. Bahlo M, Bromhead CJ. Generating linkage mapping files from Affymetrix SNP chip data. Bioinformatics 2009;25(15):1961–1962.

26. Arias M, Garcia-Murias M, Sobrido MJ. Spinocerebellar ataxia 36 (SCA36): <<Costa da Morte ataxia>>. Neurologia 2017;32(6):386–393.

27. Chao MJ, Kim KH, Shin JW, et al. Population-specific genetic modification of Huntington’s disease in Venezuela. PLoS Genet 2018;14(5):e1007274.

28. Jayadev S, Michelson S, Lipe H, Bird T. Cambodian founder effect for spinocerebellar ataxia type 3 (Machado-Joseph disease). J Neurol Sci 2006;250(1-2):110–113.

29. Valera JM, Diaz T, Petty LE, et al. Prevalence of spinocerebellar ataxia 36 in a US population. Neurol Genet 2017;3(4):e174.

30. Coutelier M, Stevanin G, Brice A. Genetic landscape remodelling in spinocerebellar ataxias: the influence of next-generation sequencing. J Neurol 2015;262(10):2382–2395.

31. Galatolo D, Tessa A, Filla A, Santorelli FM. Clinical application of next generation sequencing in hereditary spinocerebellar ataxia: increasing the diagnostic yield and broadening the ataxia-spasticity spectrum. A retrospective analysis. Neurogenetics 2018;19(1):1–8.

32. Sullivan R, Yau WY, O’Connor E, Houlden H. Spinocerebellar ataxia: an update. J Neurol 2019;266(2):533–544.

